# Multifactorial approach is needed to unravel the maturation phases of human neurons derived from induced pluripotent stem cells

**DOI:** 10.1101/2023.09.21.558836

**Authors:** Maissa Ben Mahmoud, Anikó Rátkai, Krisztina Bauer, Norbert Bencsik, Attila Szücs, Katalin Schlett, Krisztián Tárnok

## Abstract

Neurons derived from induced pluripotent stem cells (h-iPSC-Ns) provide an invaluable model for studying the physiological aspects of neuronal development and diseases. However, multiple studies have also demonstrated that h-iPSC-Ns exhibit a high degree of functional and epigenetic diversity. Due to the imprecise characterization and significant variation among the currently available maturation protocols, it is essential to establish a set of criteria to standardize models and accurately characterize and define the developmental properties of neurons derived from iPSCs.

In this study, we conducted a comprehensive analysis of the h-iPSC-Ns via electrophysiological and microscopic techniques to follow their functional development at the cellular and network levels. This enabled us to provide a thorough description of the maturation process of h-iPSC-Ns over a 10-week period *in vitro*. Specifically, we have used conventional whole-cell patch-clamp and dynamic clamp techniques, alongside morphometry, to assess the characteristics of maturing h-iPSC-Ns. Additionally, we utilized calcium imaging to monitor the progression of synaptic activity and network communication. At the single cell level, human neurons exhibited gradually decreasing membrane resistance in parallel with improved excitability by 5 weeks of maturation. Their firing profiles were consistent with those of mature regular firing type of neurons. At the network level we observed the development of abundant fast glutamatergic and depolarizing GABAergic synaptic connections together with synchronized network activity. The identified sequence of differentiation events are consistent and offers a robust framework for developing targeted experiments at varying stages of neuronal maturation. This framework allows for the use of different, age-related methodologies or a singular set of experiments for a culture’s maturation.

## Introduction

Induced pluripotent stem cell (iPSC) technology, developed by Yamanaka and Takahashi in 2006, provides a hopeful answer to the ethical and technical difficulties involved in using living human brain tissue. It also offers an opportunity to overcome the inadequacies of animal models used in neurodegenerative disease studies, which only partially recapitulate the precise mechanisms of disease development and progression, especially in late-onset diseases (Shi *et al*., 2017). iPSCs are generated *in vitro* from somatic cells, mostly from skin or blood, by ‘reprogramming’ them to an embryonic pluripotent state. This broad developmental potential allows the generation of human cells, such as neurons, for therapeutic purposes or the construction of disease models, enabling the analysis of disease aspects that cannot be studied in patients or animal models (Ardhanareeswaran *et al*., 2017). Such stem cell-based test systems have several key advantages over traditional animal models, including the ability to (i) study and characterize human-specific, disease-related cell types that model early aspects of human brain development (Yang and Shcheglovitov, 2020), (ii) reflect the genetic background of patient groups (Deneault *et al*., 2019), and (iii) be used for high-throughput screening of drug candidates (Silva and Haggarty, 2020).

Neurons derived from human iPSCs (h-iPSC-Ns) have been proven particularly useful in elucidating the complex molecular mechanisms of neurodevelopmental disorders and monogenic brain diseases (Wilson and Newell-Litwa, 2018; Autar *et al*., 2022; Sahlgren Bendtsen and Hall, 2023). Several studies have investigated the electrophysiological and morphological properties of h-iPSC-Ns (Prè *et al*., 2014; Zhang *et al*., 2017; Xie *et al*., 2018). It has been shown that h-iPSC-Ns exhibit electrical properties similar to those observed in mature neurons, such as action potentials, synaptic currents and spontaneous firing, as well as morphological maturation of cultured neurons (Autar *et al*., 2022). However, studies have also shown that iPSC derived neurons are functionally and epigenetically diverse (Aversano, Caiazza and Caiazzo, 2022). Multiple differentiation and maturation protocols of h-iPSC-Ns derived from different somatic cell types have resulted in only approximate time points (Kang et al., 2017) when neurons are regarded as ‘mature’. This is based on a successive progression of selected neuronal hallmarks (morphological, electrophysiological or gene expression characteristics) which seem to reproduce a given set of neuronal functions observed in the developing neuronal network during the early and middle stages of neuronal development. On the other hand, immature characteristics pose a challenge when using h-iPSC-Ns to study age-related pathophenotypes (Sheng *et al*., 2018; Olova *et al*., 2019). Due to the imprecise categorization and significant disparity among existing maturation protocols, standardization of the models is necessary. Therefore, it is crucial to define and establish a set of criteria for characterizing the developmental properties of neurons derived from iPSCs (Anderson et al. 2021).

In the present study, we have conducted a detailed characterization of the h-iPSC-Ns by combing individual cell- and network-level experiments allowing a profonde description of the *in vitro* maturation process of the h-iPSC-Ns over 10 weeks. Specifically, we have utilized conventional whole cell patch-clamp and dynamic clamp along with morphometry to evaluate the individual properties of the maturing h-iPSC-Ns, and calcium imaging to follow the development of the synaptic activity and network communication. All these aspects can be used as a follow-up standard of the h-iPSC-Ns maturation. The uncovered progression of differentiation events validates the usability of the model system and gives us a powerful tool to plan targeted experiments during defined stages of neuronal maturation.

## Materials and methods

### Cell culture

Human Neural Presursor Cells (NPCs, CTL1 S11, Nagy et al. 2017) were plated in Poly-L-Ornithine (PLO) (0.01%, Sigma, #P4957) and Laminin (3μg/cm^2^, Sigma, #L2020) coated 6-well dishes. The culturing media contained DMEM/F12 GlutaMAX (Gibco, #31331-028) and Neurobasal (Gibco, #21103-049) with 1% of N2 Supplement (Gibco, #17502-048), 2% of B27 Supplement (Gibco, #17504-044)), 2mM of Glutamax (Gibco, #35050061), non-essential amino acid solution (Sigma, #M7145) and antibiotic antimycotic solution (Sigma, #A5955). 10 ng/ml fibroblast growth factor basic (FGF-basic, AA 1-155, Recombinant Human Protein #PHG0264), 10 ng/ml epithelial growth factor (EGF, Recombinant Human Protein #PHG0311) and 10 µM ROCK Inhibitor Y-27632 (Sigma, #Y0503) were added to the media freshly before use. The cells were grown to confluence, then passaged after six or seven days. For terminal differentiation, 10^5^ cells / well were plated into 24-well plates containing PLO-Laminin coated glass coverslips. After 24 hours, culturing medium was changed to the Differentiation Medium which contained BrainPhys Neuronal Medium (Stem Cell Technologies #05790) with 1% of N2 Supplement (Gibco, #17502-048), 2% of B27 Supplement (Gibco, #17504-044), 1 µg/ml Laminin (L2020). Growth factors (10 ng/ml brain derived neurotrophic factors (BDNF, Gibco, #PHC7074); 10 ng/ml Glial derived neurotrophic factor (GDNF, Gibco, #PHC7045); 1 mM Dibutyryl cyclic-AMP (Sigma, #D0627); 200 nM ascorbic acid (Sigma, #A8960) were added freshly to the medium and a half media change was performed twice a week during the differentiation.

### Sholl analysis

Biocytin-filled recorded cells from week 1-7 were fixed with 4% paraformaldehyde (in phosphate-based saline (PBS), pH 7.4) for 20 minutes, visualized with streptavidin-TRITC (1:500, #016-020-084, Jackson ImmunoResearch) and mounted in Mowiol 4.88 with DAPI (Polysciences). Cells were imaged with a Zeiss CellObserver Z1 widefield fluorescent microscope, using Plan-Apochromat 20×/0.8 (Zeiss) objective and the images were acquired as z-stack tiles with Axiocam MRm camera (Zeiss) using the ZEN software (version 2.6, Zeiss). On the microscopic images, neuronal shape was traced by the Simple Neurite Tracer plugin (Arshadi *et al*., 2021) with Fiji (ImajeJ framework) software (Schindelin *et al*., 2012) (https://imagej.net/plugins/snt/). Using this plugin, a line stack of the traces was created for subsequent Sholl analysis, performed using the Sholl analysis plugin (https://imagej.net/plugins/sholl-analysis) with a starting radius of 10 µm and final radius at the furthest point of the cell in increments of 10 µm. Data relating to neurite length and intersection were obtained using the simple neurite tracer plugin. Process length data presented as box plots of interquartile range together with individual data points. Normalized branching data (intersections per µm) are presented as mean ± SE.

### Electrophysiology

#### Patch clamp recordings

Electrophysiological recordings were performed under an Axiovert 200M microscope (Zeiss) equipped with EC Plan-Neofluar 20×/0.5 objective. Spontaneous synaptic activity and evoked responses were recorded at room temperature (21-23 °C) in whole-cell conditions using a MultiClamp 700B amplifier (Molecular Devices). Intracellular voltage and current traces were sampled at 20 kHz and stimulus command waveforms were generated by the data acquisition software DASYLab v.11 (National Instruments). Patch pipettes (7-10 MOhm) were pulled from standard wall glass of 1.5 mm OD (World Precision Instruments). The composition of the bath solution (ACSF) was (in mM): NaCl 140, KCl 5, CaCl_2_ 2, MgCl_2_ 1, HEPES 5, D-glucose 10; pH set to 7.45, while patch electrodes were filled with the following solution (in mM): K-gluconate 100, KCl 10, KOH 20, MgCl_2_ 2, NaCl 2, HEPES 10, EGTA 0.2, D-glucose 5; pH set to 7.3. For subsequent morphological analysis, 0.5 mM biocytin (Tocris, #33-491-0) was added to the patch pipette solution. To record voltage responses of the neurons and to extract multiple physiological parameters of those, we used a current step stimulation from week 1 to week 10. Stepwise current commands of 350 ms duration, starting at – 20 pA and incremented by +2 pA were delivered in 1.25 s cycles. In such experiments, multiple physiological parameters including the input resistance, membrane time constant, spike amplitude and half-width were determined for each cell.

Spontaneous excitatory postsynaptic currents (sEPSCs) were acquired at − 60 mV holding potential in voltage clamp mode. To characterize the chemical properties of the postsynaptic currents, selective pharmacological blocking of AMPA-, NMDA- and GABA_A_- receptors was carried out by using CNQX (10 μM, Tocris), AP-5 (40 μM, Tocris) and bicuculline (30 μM, Tocris), respectively. Analysis of both the sEPSCs and the evoked responses was performed using a software developed by A. Szücs (NeuroExpress), (https://www.researchgate.net/publication/323547374_Analysis_of_miniature_excitatory_postsynaptic_currents_mini_analysis_in_NeuroExpress_Program_available_for_download).

### Dynamic clamp experiments

We used the dynamic clamp technique to investigate the neurons’ firing responses under physiologically more realistic inputs. Here, we exposed the neurons to simulated oscillatory synaptic inputs consisting of one excitatory and one inhibitory channel (Szücs and Huerta, 2015). The frequency of the oscillatory envelope of the excitatory synaptic input was either 4 Hz (*low frequency* stimulation) or 20 Hz (*high frequency* stimulation). Stimulus waveforms serving as presynaptic voltage for the dynamic clamp system were generated by the data acquisition software DASYLab v.11 (National Instruments). To compute the synaptic conductance and the net postsynaptic current we used the StdpC v.2012 program (Nowotny *et al*., 2006). The synaptic time constant was 10 ms for both the excitatory and inhibitory connections, their reversal potential was 0 mV and -72 mV, respectively. The excitatory synaptic conductance (matching that of the value of the inhibitory conductance) was set in a way that approximately 10 action potentials were evoked under a single sweep of the oscillatory theta waveform.

### Calcium imaging

h-iPSC-Ns cultivated on glass cover coverslips at 4, 6 and 8 weeks of maturation were loaded with 2.5 µM Fluo-3-AM (Molecular Probes, #F1242) dissolved in differentiation medium for 30 min at 37 °C and at 5% CO_2_. Subsequently the differentiation medium was changed to BrainPhys™ Imaging Optimized Medium (StemCell technologies, #05796) for image acquisition. Imaging was conducted using a Spinning Disc confocal system (Zeiss), in an environmental chamber at 37 °C and 5% CO_2_. Live cell imaging of the calcium spatial dynamics was detected by exciting the Fluo-3-AM with a combination of excitation laser wavelength at both 488 and 514 nm. Images were captured for 5min with 500 ms frame rate using a Zeiss Axio Observer SD spinning disc confocal microscope equipped with a LCI Plan-Apochromat 25x/0.8 oil objective at 520-560 nm. Data were analyzed using the Mesmerize software (Kolar et al., 2021). Somatic regions of interest (ROI) were defined over every cell within the field of view. Mean fluorescent intensities of the ROIs are expressed as ΔF/F0 ratio, which represents the change in fluorescent intensity (ΔF) normalized to the baseline fluorescence of Fluo-3 (F0). Based on the calcium transients (ΔF/F0), heatmaps were generated, where results are depicted using a standard z score (0-6) and a color scale, where 0 denotes the lack of a Ca-transient and 6 represents the maximum relative intensity of calcium transients.

To detect synchronization of Ca-transients as indicatives of the development of robust synaptic connectivity, we calculated pairwise Pearson correlations for the ROI channels. Here, first we established the sequences of the arrival times of Ca-waves and created event density functions by convolving these arrival times with a Gaussian function (kernel, 5 s half-width). The correlation coefficients for such smooth and continuous density functions were calculated and plotted in the form of the correlation matrix. Averaged values obtained from independent experiments were used to calculate the interevent interval, peak halfwidth and Spearman correlation of the mentioned independent variable.

### Quantitative real-time PCR (qRT-PCR)

Cultures were lysed and RNA samples were obtained using Quick-RNA MiniPrep (ZYMO Research, USA). Total RNA was quantified with a spectrophotometer (Implen, NanoPhotometer N60). Reverse transcription was performed with the Maxima First Strand cDNA Synthesis for RT-qPCR kit (Thermo Scientific) according to the manufacturer’s instruction. Messenger RNA expression was investigated using the Maxima SYBR qPCR Master Mix (Thermo Scientific) with specific primers (**Table 1**). The qPCR run was performed using a CFX96 (C1000 Touch) from Bio-Rad Laboratories (USA) with the following settings: 1 cycle at 95 °C for 10 min, 40 cycles at 95 °C for 15 sec followed by 55 °C for 30 sec and 72 °C for 30 sec. Cq and RFU values were obtained from Bio-Rad CFX Manager software (Bio-Rad, USA). Cq values were normalized to housekeeping genes GAPDH and RLP-13a. The relative expression of the given genes was calculated using the ΔΔCq method from triplicated samples using the CFX Manager software (version 3.1, BioRad).

**Table 1.**
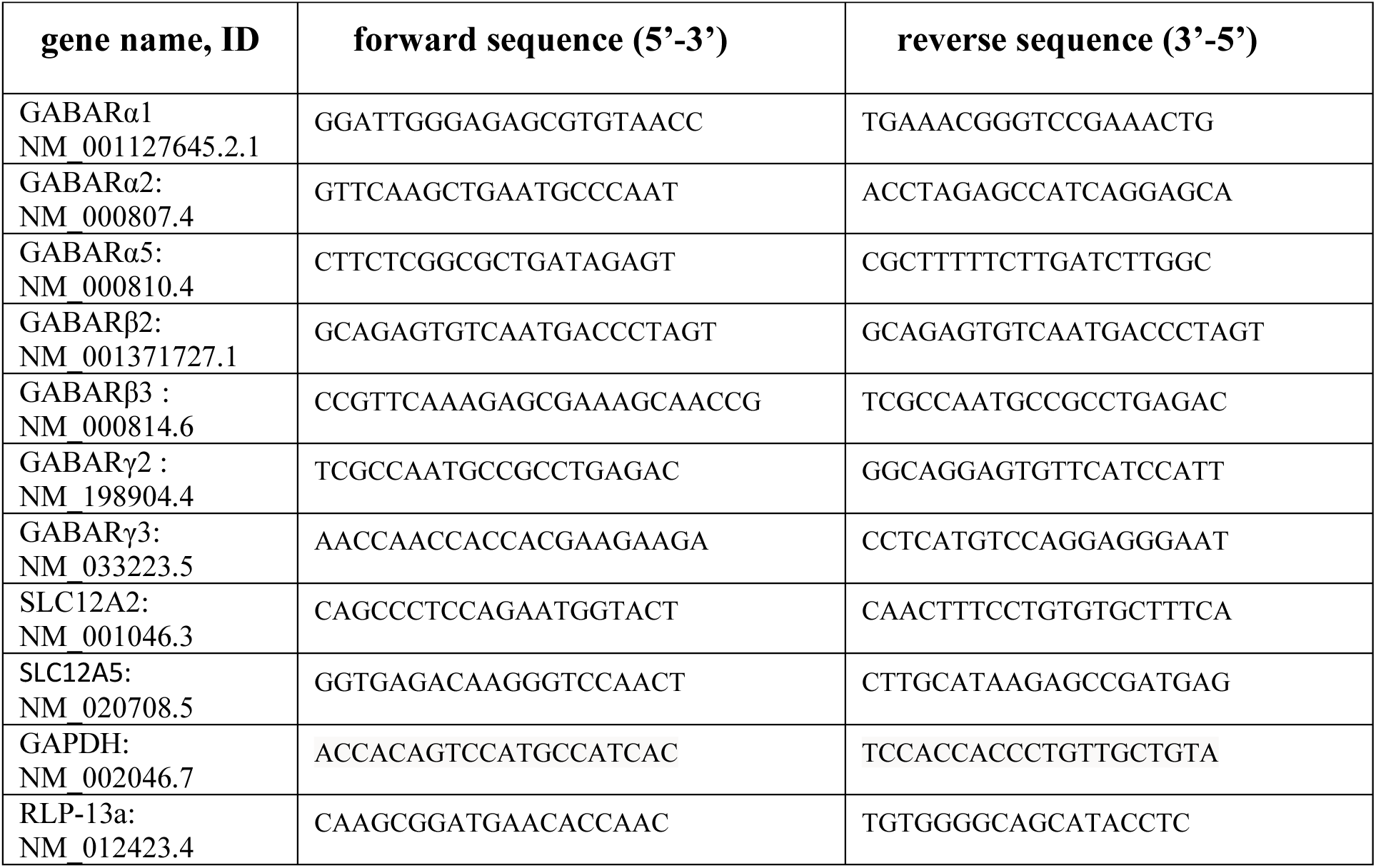
Primer sequences used in RT-qPCR experiments.

### Statistical analyses of the data

One-way ANOVA, Student’s t test or non-parametric Mann-Whitney tests were used for statistical evaluation unless otherwise indicated. SPSS Statistics (IBM) was used to calculate statistics. Data are displayed as mean ± SEM, unless otherwise indicated. p values were accepted statistically significant as * p<0.05, ** p<0.01, *** p<0.001.

## Results

### Morphological maturation of human iPSC-derived neurons is enhanced after 5 weeks

To track the development and maturation of human cortical neurons *in vitro*, we used a partially characterized induced pluripotent stem cell (iPSC) line (CTL1 S11) originated from a neurotypic, young adult male person (Nagy *et al*., 2017). The S11 cells can be differentiated into forebrain neurons and astroglia cells by the 2-inhibitor method (Nagy *et al*., 2017). In our experiments, human neurons were differentiated from S11 h-iPSC-derived neuronal precursor cells (S11 NPCs) in neuronal differentiation media supplemented with BDNF (10 ng/mL), GDNF (10 ng/mL) and dibutyryl-cAMP (1 mM) and were cultured for 10 weeks, *in vitro*.

Neuronal process-formation (**Fig. 1A-D**) and the presence of neuronal markers (IIIβ-tubulin, MAP2, data not shown) were detected from the first week in culture, together with the first appearance of astroglia cells (data not shown). We carried out whole-cell patch clamp measurements where cells were filled up with biocytin after the electrophysiological recordings to assess the morphometric characteristics of the developing neurons parallel to their physiological aspects (see the recorded traces below the corresponding images). Based on the fluorescent immunocytochemistry visualization of the biocytin signal, we reconstructed and analyzed the cell morphology within a 350 µm radius from the soma (**Fig. 1**). Sholl analyses indicate that both the total dendritic length and complexity of the arborization increase in a biphasic manner (**Fig. 1E, F**). The initial process expansion was slowly followed by the branching of processes, however after the 5^th^ week of culture, elongation and branching accelerated, leading to more elaborate morphology (**Fig. 1D**) together with the appearance of mature electrophysiological properties in the neurons (**Fig. 2**).

**Figure 1.**
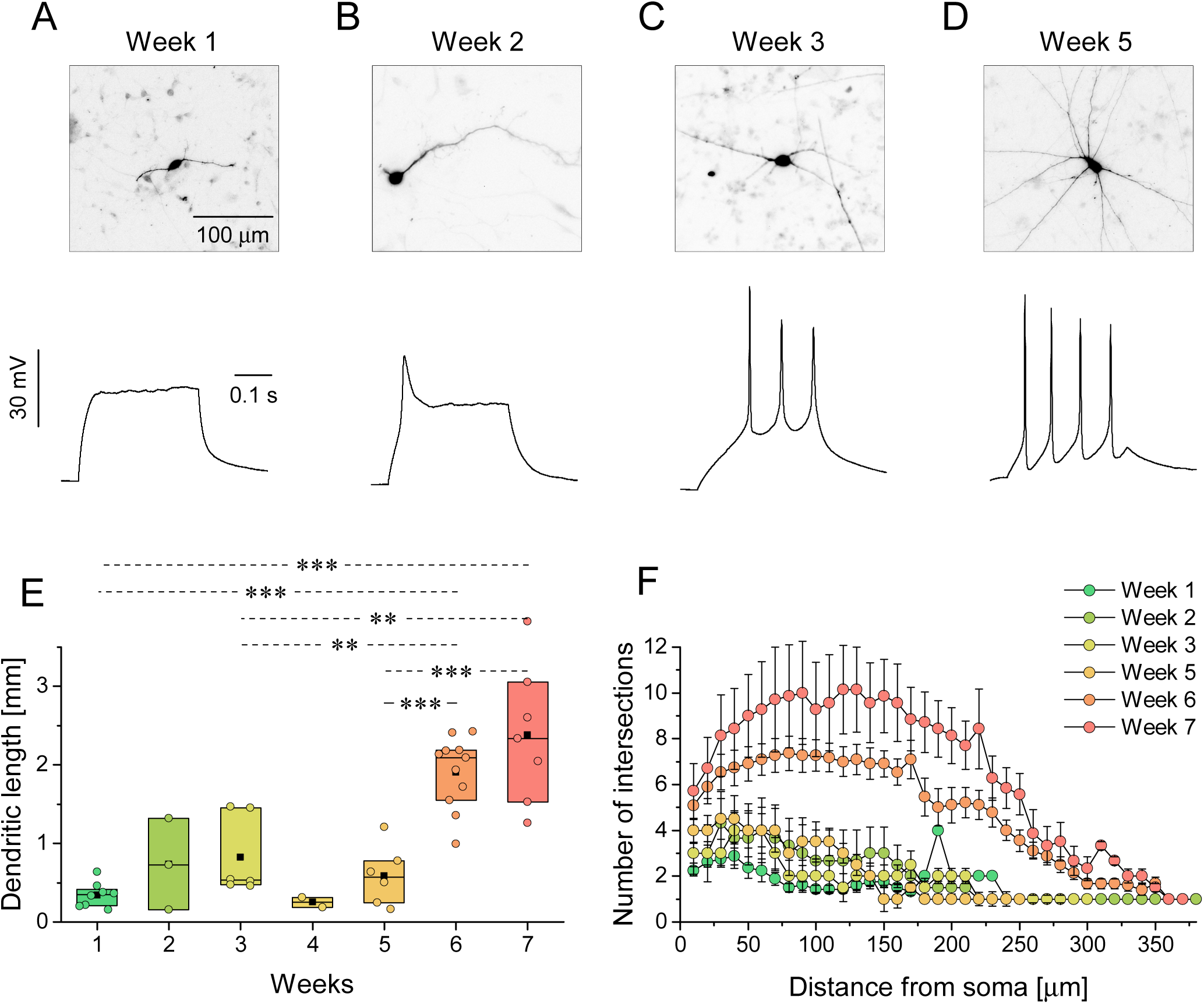
Electrophysiological and morphological properties develop in a time-dependent manner. (**A-D**) Cells filled with biocytin during patch clamp recordings were used for the morphological study. Voltage responses were recorded using standard current clamp protocols under whole-cell condition. The arborization of the biocytin+ cells was characterized by Sholl analysis, displaying (**E**) the total dendritic length and (**F**) the arborization of neural processes represented as the number of intersections in the function of the distance from the soma. Data are presented as box plots of interquartile range together with individual data points (n = 8, 3, 6, 2, 6, 11, 7 for week 1-7 respectively). ***: p < 0.001, **: p < 0.01, *:p < 0.05

**Figure 2.**
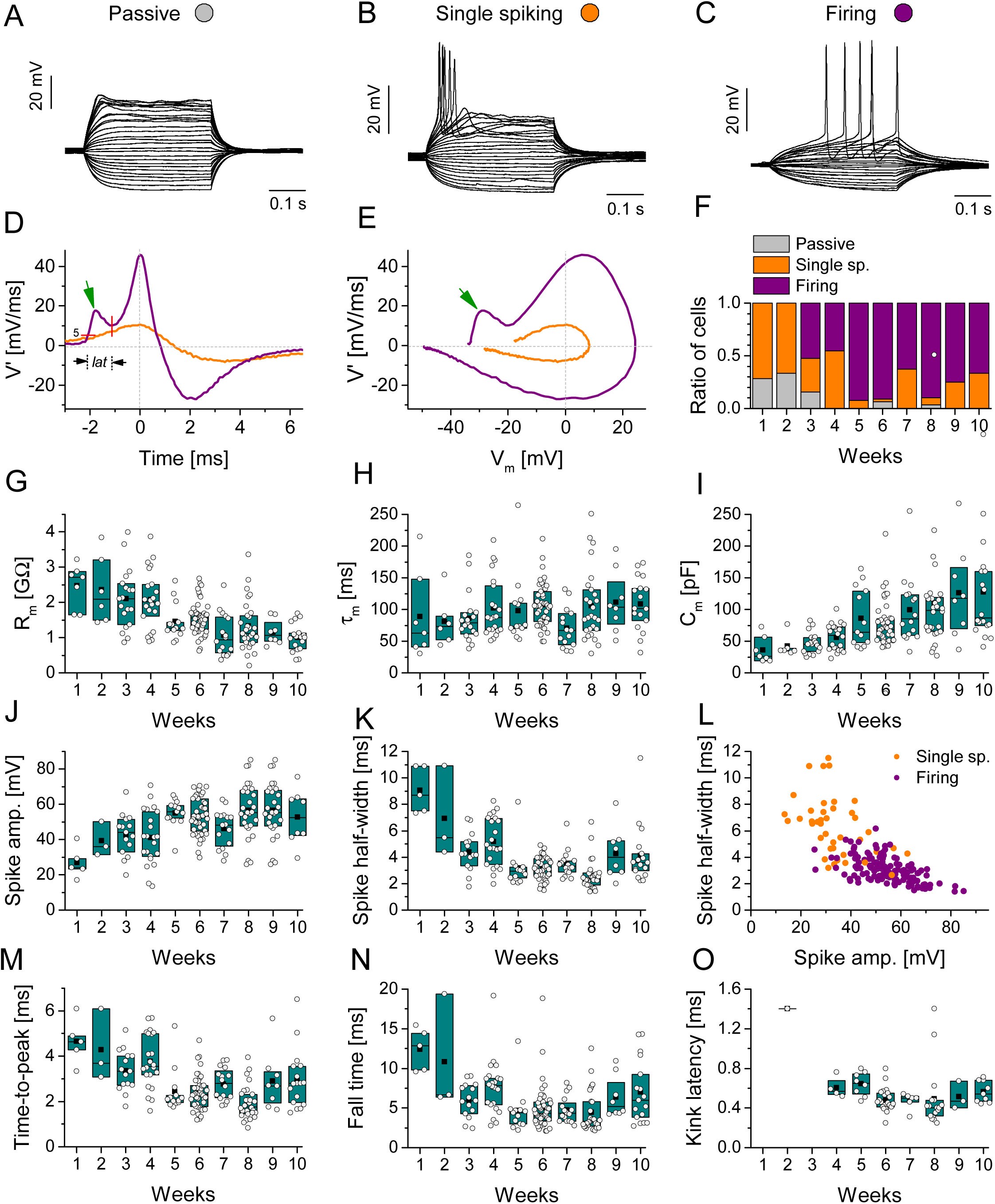
Evolution of passive and active membrane properties during h-iPSC-Ns maturation. (**A-C**) Three distinct electrophysiological phenotypes - passive (**A**), single spiking (**B**) and firing (**C**) - were observed under current clamp protocols in whole cell condition during maturation. (**D**) First derivative of the membrane potential and the phase portrait (**E**) differ between single spiking (orange) vs. firing (purple) phenotypes. The green arrows indicate the prominent transient fluctuation of the AP slope (*kink*) in the firing phenotype. (**F**) Ratios of the three electrophysiological phenotypes over time during 10 weeks of maturation. (**G-I**) Passive membrane properties during 10 weeks of maturation. (**G**) Average membrane resistance (Rm) decreased over time. (**H and I**) Membrane time constant (τm) and the membrane capacitance (Cm) increased during the 10 weeks of maturation. (**J, K, M, N and O**) Active membrane properties during the 10 weeks of maturation. (**J**) Spike amplitude of action potentials showed a time dependent increase with a stabilization phase around the 5 weeks of maturation. Spike half-width (**K**), time-to-peak (**M**) and fall time (**N**) decreased in a time-dependent manner, respectively. (**L**) Negative correlation between the spike amplitude and half-width of the firing phenotypes is observed. (**O**) Kinks can be detected, and kink latency values can be calculated for the firing type h-iPSC-Ns from the 4th week of maturation. Data presented as box plots of interquartile range together with individual data points. The total number of cells examined for these experiments was n=108.

### Multifactorial analysis reveals the timescale of stabilizing neuronal electrophysiological properties during maturation

To determine the physiological maturation stages of human iPSC-derived neurons over time, patch clamp measurements in whole-cell configuration were performed weekly from the 1^st^ week (DIV8, days in vitro) to the 10^th^ week (DIV74). Our experiments focused on the analysis of both intrinsic biophysical properties and the excitability of h-iPSC-Ns. Multiple passive and active physiological parameters were calculated from the voltage responses of neurons under incrementing levels of the injected current (**Fig. 2A-C**). The physiological parameters and intrinsic excitability of h-iPSC-Ns were analyzed as functions of the maturation time (**Fig. 2G-O**). Initially, most h-iPSC-Ns exhibited passive (resistive-capacitive-type) membrane response (**Fig. 2A and F**) indicating the lack of voltage-dependent membrane currents. The average membrane resistance was at maximal level in the first week of maturation and gradually decreased over time (**Fig. 2G**). The membrane time constant (ρ_m_) remained fairly consistent across our experiments (**Fig. 2H**), however, we observed a monotonous increase of the calculated membrane capacitance during the 10 weeks (**Fig. 2I**). This observation agrees with the morphological maturation of the cells, i.e. the development of extended processes and increase of total membrane surface. From the 3^rd^ week, h-iPSC-Ns displayed action potentials first with single spike responses (**Fig. 2B**) and later with repetitive firing (**Fig. 2C**) under depolarizing current steps. Based on the propensity of neurons to fire action potentials we established three functional phenotypes, namely passive, single spiking and firing neurons (**Fig 2A-C**).

The percentage of firing type neurons increased over time (**Fig. 2F)**. Action potentials became gradually more pronounced and sharper as revealed by the analysis of the first-time derivative of the membrane potential (**Fig. 2D and E**). Single spiking neurons typically emitted action potentials with slow dynamics (slope remaining below 20 mV/ms; see the orange curves). The phase portrait on **Fig. 2E** also demonstrates the higher amplitude and faster dynamics of the action potentials in the case of firing phenotype (see the purple curves). We also noticed the high abundance of firing responses with a characteristic bump (also referred to as *kink*) in the phase portraits (**Fig. 2D and E**, green arrow). This feature indicates a temporal mismatch between the activation of transient Na-currents in the soma vs. axon initial segment (Bean, 2007). By detecting the onset and termination of the kink in the action potential waveforms (**Fig. 2D**, *lat*) we extracted a new parameter called kink latency and found that it was fairly stable across our measurements after 4 weeks *in vitro* (**Fig. 2O**). Our patch clamp measurements also revealed a time-dependent increase of the spike amplitude (**Fig. 2J**) and decrease in the half-width of action potentials (**Fig. 2K**). Here, steep changes were observed until cca. 5 weeks of maturation and then parameters stabilized with less variability. Spike rise times (time-to-peak, **Fig. 2M**) and spike fall times (**Fig. 2N**) exhibited a similar biphasic change over the time of the study.

Interestingly, by examining the correlation between spike amplitude and half-width we found a clear separation of single spiking vs. firing neurons, where members of the two phenotypes formed distinct clusters (**Fig. 2L**).

### iPSC-Ns fire reliably under simulated synaptic inputs from the 4^th^ week of maturation

To further characterize the excitability and integrative properties of h-iPSC-Ns, we exposed them to simulated synaptic inputs via dynamic clamp (Szabó *et al*., 2021). In this configuration, we can mimic *in vivo* conditions (temporally complex synaptic inputs with amplitude fluctuations) resembling to what inputs neurons are exposed in forming brain circuits. To achieve this, we first designed excitatory and inhibitory conductance waveforms consisting of variable amplitude synaptic transients (**Fig. 3A** and **C**, g_exc_ and g_inh_). The excitatory input (g_exc_) was used to deliver oscillatory drive to the neurons either as a low frequency input (4 Hz) or a high frequency rhythm (20 Hz). The inhibitory input (g_inh_) was non-periodic and served as a random inhibitory background. We set the maximal synaptic conductance of the excitatory input in a way that neurons emitted cca. 10 spikes under one sweep of the theta stimulation. Next, we repeated the stimulation 20 times for both the 4 Hz and 20 Hz input and observed the spike responses. Under such scenario, neurons tend to fire their action potentials in well-defined locations along the stimulus (time windows) and these can be referred to as spike events. The reliability and temporal precision in such spike events can be then calculated.

**Figure 3.**
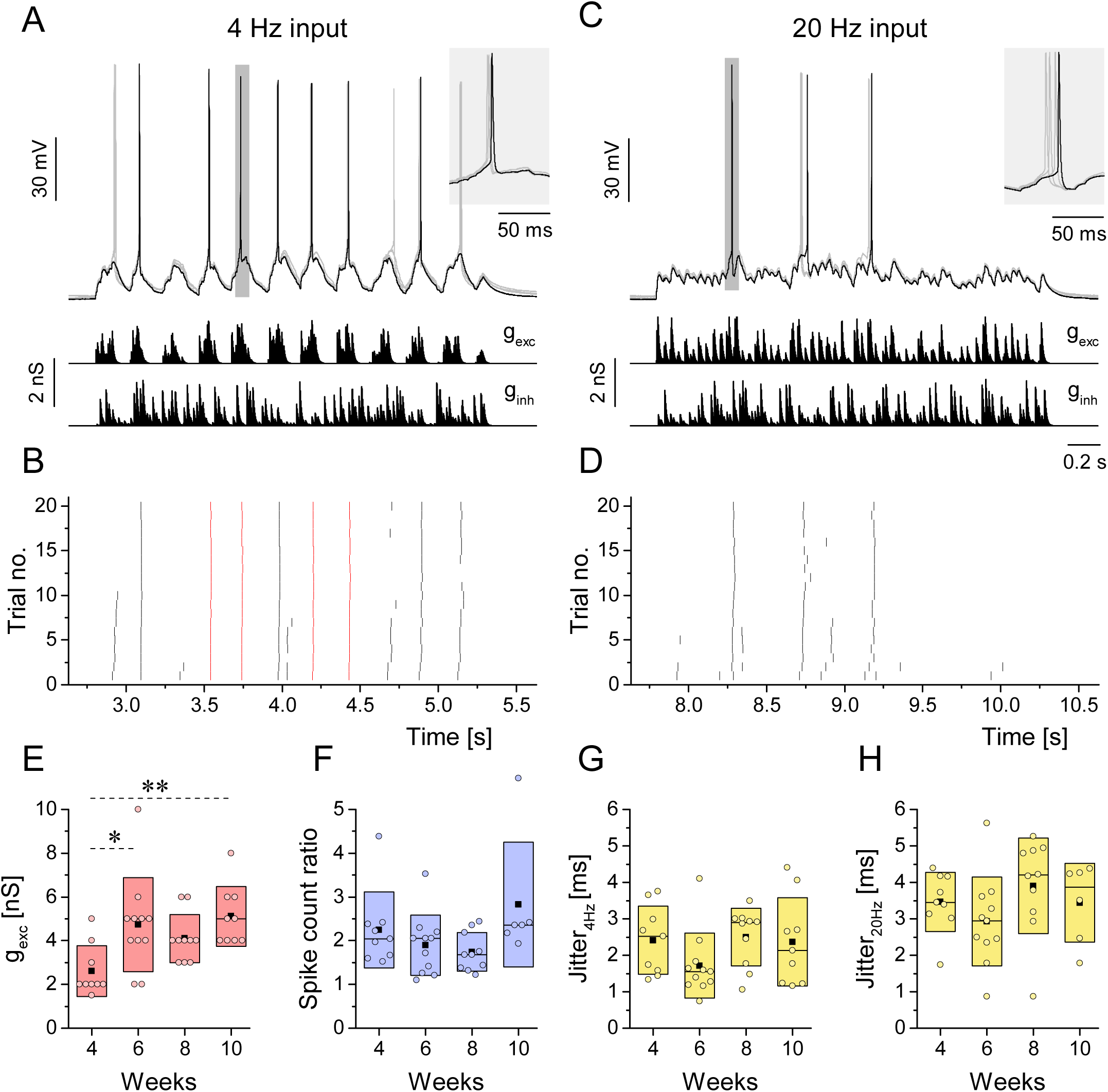
Effects of simulated oscillatory synaptic inputs on h-iPSC-Ns under dynamic clamp condition. (**A**) Four overlapping voltage traces of the neurons under simulated 4Hz oscillation. Conductance waveforms of the excitatory (gexc) and the inhibitory (ginh) inputs are shown below the traces. (**B**) Peri-stimulus spike raster plot of the firing response of the neurons under 4Hz oscillation. Red spikes indicate events with spike jitter less than 2 ms. (**C**) Four overlapping voltage traces of the neurons under simulated 20 Hz inputs. (**D**) Corresponding peri-stimulus spike raster plot for the 20 Hz oscillation. (**E**) Excitatory synaptic conductance required for the emission of cca. 10 spikes under the 4Hz inputs (inhibitory synaptic strength matches that of the excitation). More mature h-iPSC-Ns required stronger excitatory or inhibitory conductance to fire the targeted spike number. (**F**) In average, twice as many spikes were evoked by the 4Hz input than by the 20 Hz input (4Hz /20 Hz spike count ratio). These ratios were fairly stable during the development (4, 6, 8 and 10 weeks). (**G** and **H**) Spike jitter upon 4Hz (**G**) and 20 Hz (**H**) inputs indicate higher timing precision in case of lower frequency stimuli. Data presented as box plots of interquartile range together with individual data points. ***: p < 0.001, **: p < 0.01, *:p < 0.05

In the example of **Fig. 3A**, the neuron emitted its action potentials on the top of the excitatory waves and did it in a reliable fashion. Peri-stimulus spike raster plot of this experiment is shown in **Fig. 3B**. Ticks with red color indicate the spike events where the temporal jitter of spikes was under 2 ms. Comparing the raster plots of the low vs. high frequency stimuli we find far less spikes and higher temporal jitter in the latter case (**Fig. 3D**). Consequently, h-iPSC-Ns exhibit lower (4 Hz) frequency-preference in their firing output, i.e. they fire more spikes and with higher precision than when receiving high-frequency signals. The maximal synaptic conductance used for these experiments is shown in **Fig. 3E**. Reaching ten spikes per sweep required higher values of g_exc_ as the neurons matured. This finding well agreed with the fact that the membrane resistance of the neurons tended to drop in the same time period (**Fig. 2G**). All the neurons in this experiment exhibited cca. 2 times more spikes under 4 Hz than 20 Hz inputs as shown by the low/high frequency ratio plot (**Fig. 3F**) independently from the *in vitro* age. Spike jitter calculated from firing responses under low and high frequency inputs did not change significantly during the maturation (**Fig. 3G** and **H**).

### Fast excitatory AMPA- and slower depolarizing GABA-inputs are abundant in the developing h-iPSC-Ns

In our following experiments, we investigated the development of chemical synaptic connections in the networks of h-iPSC-Ns. Voltage clamp and selective blockers of putative postsynaptic receptors of glutamatergic and GABAergic neurotransmission were used to reveal the properties of synaptic communication between the developing neurons. Typically, we observed sEPSCs that exhibited either fast (decaying within 10 ms) or slower (decay >50 ms) kinetics (**Fig. 4A, C, E**). CNQX (10 μM), a selective blocker of the AMPA receptors, reliably eliminated the fast kinetics events while not affecting the slower ones (**Fig 4B**). To check whether NMDA-receptors were contributing to the observed postsynaptic currents, we used AP-5 (40 μM) but found no changes in the appearance of the sEPSCs (**Fig. 4D**). To identify the source of the slower sEPSCs we used bicuculline (30 μM), a GABA_A_-receptor antagonist. This treatment completely eliminated the slow events, but the fast AMPA-events were not affected (**Fig. 4F**). We also noted that the estimated reversal potential of the slower currents (cca. -25 mV, not shown) was more hyperpolarized than that of the AMPA-events, still, these postsynaptic currents impose a strong excitatory drive on the developing h-iPSC-Ns. Therefore, slow kinetics sEPSCs were generated by excitatory GABA transmission mimicking the early steps of neuronal development (Hutcheon *et al*., 2000). In contrast, the fast sEPSCs have the characteristics of typical AMPA-type glutamatergic postsynaptic currents in mature neurons.

**Figure 4.**
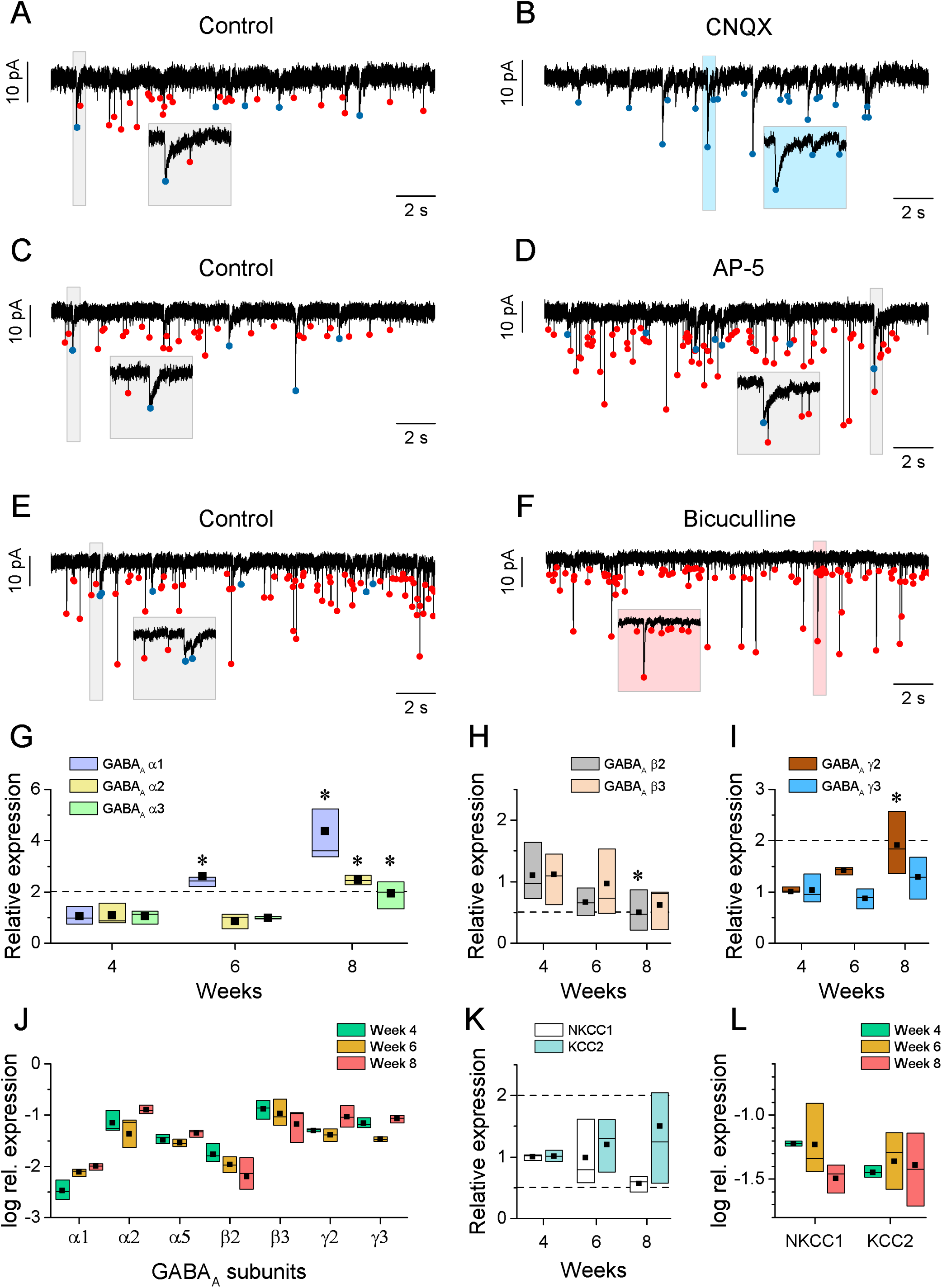
Excitatory fast glutamatergic and slow GABAergic synaptic EPSCs in h-iPSC-Ns are detected at 6^th^ week of maturation. (**A, C and E**) Slow (blue dots) and fast (red dots) spontaneous postsynaptic currents (sEPSCs) were detected in control condition. (**B**) CNQX (10 μM), a selective competitive AMPA/Kainate receptor antagonist blocked the fast sEPSCs. (**D**) Fast and slow sEPSCs persist after application of a selective NMDA antagonist AP-5 (40 μM). (**F**) Bicuculline (30 μM), a competitive GABAA receptor antagonist blocked the slow sEPSCs without affecting the fast events. **GABAA subunit expression changes in a time-dependent manner**. (**G**) Relative expression of the GABAA *alpha* 1, *alpha* 2 and *alpha* 5 subunits (**H**), GABAA *beta* 2 and *beta* 3 subunits (**I**) and GABAA *gamma* 2 and *gamma* 3 subunits (**K**) at 4, 6 and 8 weeks of maturation normalized to gene expression level at 4 weeks. To show comparable expression of different subunits, GABAA subunit values were normalized to the expression of human ribosomal protein L13a (*RLP13a*) at 4, 6 and 8 weeks of maturation (**K**) and expressed as log rel. expression. Relative gene expression of the *NKCC1* transporter and *KCC2* symporter at 4, 6 and 8 weeks of maturation normalized to the 4week data (**L**). *NKCC1* transporter and *KCC2* symporter normalized to the logarithmic expression of the human ribosomal protein L13a at 4, 6 and 8 weeks of maturation. Data presented as box plots of interquartile range. *: p < 0.05 (One-way ANOVA, Tukey post-hoc test).

Based on the slow sEPSCs observed by electrophysiology, we decided to investigate the changes in the gene expression levels of commonly occurring GABA_A_ ionotropic receptor subunits *(alpha 1, 2, 5; beta 2, 3; gamma 2, 3*) by qRT-PCR during the maturation of h-iPSC-Ns. Based on our results, all types of the investigated subunits were expressed in h-iPSC-Ns regardless of the developmental age (**Fig. 4J**). However, subunit-specific differences were found in the relative expression data compared to those of week 4. The relative expression of *alpha* subunits increased over time, especially the amount of *alpha 1* subunit showed a time-dependent change (**Fig. 4G**). For *beta* subunits, the expression level of both *beta 2* and *beta 3* decreased during maturation (**Fig. 4H**). In the case of the *gamma 2* subunit, an age-dependent increase was also observed **(Fig. 4I**), however the level of *gamma 3* subunits was not changed over the study period (**Fig. 4I**).

We also compared how the expression of the subunits varied relative to each other. In this case, the expression levels were normalized and plotted relative to that of human ribosomal protein L13a (*RLP13a*). *Alpha 2*, *beta 3* and *gamma 2/3* subunits were the most dominant in our cultures (**Fig. 4J**). The depolarization inducing effect of GABA_A_ receptors is due to the function of chloride transporters, therefore the expression changes of *NKCC1* and *KCC2* chloride transporters were also examined. There is a slight decrease in the level of *NKCC1* compared to 4 weeks, while there is no change in expression of *KCC2* (**Fig. 4K**), their expression levels remain similar throughout the investigated time points (**Fig. 4L**).

### Synchronous network activity follows the stabilization of electrophysiological phenotype with a latency

To evaluate the neuronal circuit forming capacities of h-iPSC-Ns over time, we recorded spontaneous calcium signals during neuronal maturation. Neurons were loaded with Fluo-3-AM dye and neuronal activity was recorded at 4, 6 and 8 weeks. Relative changes of the Ca^2+^ level within the soma of individual neurons were plotted as heatmaps (**Fig. 5A, C and E**) or normalized as correlation matrices (**Fig. 5B, D and F**). Calcium transients at 4 or 6 weeks cultures showed non-synchronized occurrences revealing a randomized neuronal activity (**Fig. 5A-D**) whereas a gradual development of h-iPSC-Ns network activity was observed, and neuronal activity was partially synchronized in 8-week-old cultures (**Fig 5E, F**). The frequency of calcium waves remained relatively stable over 4-6 weeks, but the significant decrease in the interevent intervals (**Fig. 5G**) indicated the appearance of recurring Ca-wave multiples (bursts). Concurrently, we observed the decrease of the event half-width parameter (**Fig. 5H**) that is consistent with the development of faster decaying Ca-signals. Our correlation analysis also revealed the development of network synchronization in cultures by 8 weeks of maturation (**Fig. 5E and F**). Indeed, nearly half of the ROI channels displayed recurring Ca-waves that indicated the formation of synaptically interconnected neuronal circuits. A Spearman correlation was conducted between the halfwidth of the calcium transient and the interevent interval, yielding a time-independent correlation coefficient of 0.78. This indicates that the two parameters vary in parallel over time (**Fig. 5I**).

**Figure 5.**
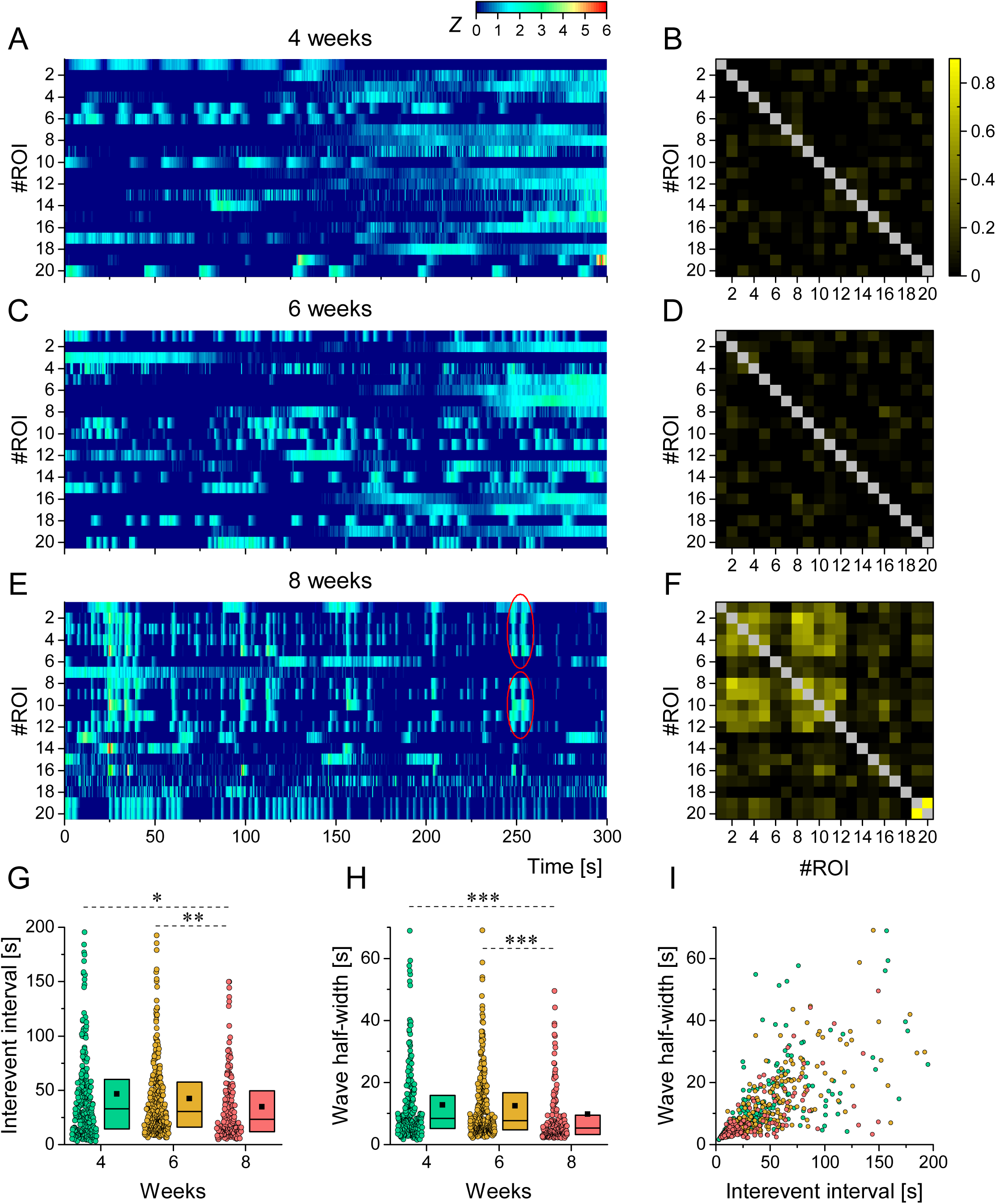
Spontaneous Ca^2+^-transients increase in h-iPSC-Ns during maturation. (A, C and E) Representative heatmaps of the Fluo-3 -AM spontaneous Ca^2+^- transients at 4, 6 and 8 weeks of h-iPSC-Ns maturation, respectively. Data from a single microscopic view consisting of 20 Regions of Interest (ROIs) are visualized using the ΔF/F0 ratio, which represents the change in fluorescent intensity (ΔF) normalized to the baseline fluorescence of Fluo-3 (F0). Synchronized Ca^2+^-signals in a few regions of interest (ROIs) are encircled in **E**. The results are depicted using a standard z score (0-6) and a color scale, where 0 denotes the lack of a Ca^2+^-transient and 6 represents the maximum of the provided collection of calcium transients. (**A**) The network exhibits slow and broad transients, consistent with early developing neurons. (**C**) Ca^2+^-transients in the neuronal network take on a more spike-like morphology reflecting what has been reported for maturing neurons. (**E**) Synchronous-like activity starts developing at 8 weeks. (**B, D and F**) Cross-correlations between the activities of the same set of 20 regions of interest (ROIs) are displayed in a symmetric matrix. Colors vary based on the Spearman correlation, where a value of 0 indicates non-correlation and 1 indicates high correlation. (**G**) The interevent interval of calcium transients was measured over a period of 4, 6, and 8 weeks during the maturation of h-iPSC-Ns. The median interevent interval of calcium transients gradually decreased during maturation. Additionally, calcium transient halfwidth (wave halfwidth) exhibited similar behavior (**H**). We established a Spearman correlation between the calcium transient halfwidth and interevent interval (**I**), revealing a time-independent correlation of 0.78. (n= 232 for 4 weeks, n=280 for 6 weeks and n=147 for 8 weeks). Data were acquired from 3 parallel cultures.

## Discussion

The application of neurons derived from human induced pluripotent stem cells (h-iPSC-Ns) as a reliable model for neuronal differentiation and drug development has been confirmed by numerous studies (Ardhanareeswaran *et al*., 2017; Silva and Haggarty, 2020; Nicholson *et al*., 2022). Since the introduction of the Yamanaka protocol (Takahashi and Yamanaka, 2006), various combinations of reprogramming, differentiation and maturation protocols have been developed to direct the reprogrammed cells towards a reliable and robust neuronal phenotype, therefore there is a need to facilitate and track neuronal differentiation via standardized methods to reduce maturation variability/heterogeneity of induced neurons (reviewed by Berry *et al*., 2019). This can be accomplished by establishing a framework of parameters to identify neuronal developmental traits, with specific emphasis on cell-autonomous and network attributes. Such guidelines can provide a valuable tool to help the establishment of targeted experiments (e.g. drug discovery) that are adapted to specific stages of neural maturation.

In the present study, we monitored the maturation of h-iPSC-Ns for up to 10 weeks by combining multiple analyses: morphometric essay, patch clamp using conventional current step protocols, dynamic clamp, Ca-imaging, and by monitoring expression changes of GABA_A_ receptor subunits and chloride transporters by qRT-PCR. Some changes of the investigated parameters suggest linear progress of maturation (such as passive membrane or network properties or network), whereas other parameters showed biphasic evolution (morphometric maturation or active membrane properties) or no development (excitability responsiveness). In addition, the phasic properties partially overlapped with each other in time.

In our timeframe, we initially observed the development of electrophysiological properties occurring in parallel with the less branched elongation of neuronal processes. There was a linear progression in the increasing conductance and decreasing membrane resistance of human neurons. On the other hand, the active membrane parameters such as the half-width and amplitude of action potentials showed the strengthening of excitability properties as a maturation-stabilization curve with an inflection point at week 5. We also notice that h-iPSC-Ns in our experiments behaved very consistently in respect to their overall cellular properties. While the variability of membrane resistance and spike shape parameters was significant, especially in early measurements, the cells exhibited typical features of regular firing neurons. In the successive period (week 6-10), firing profiles remained stable. Interestingly, the presence of the *kink* in the phase portraits of action potentials preceded the stabilization of active parameters, but was prominent and parallel with the firing phenotype from week 5. It is noteworthy that more intricate physiological features such as inward rectification, post-inhibitory rebound and burstiness were not observed in our patch clamp recordings. This indicates the lack or low level of certain voltage-dependent currents such as the hyperpolarization-activated cation current (I_h_), the low threshold Ca-current (I_T_) (Hernáth *et al*., 2019). Our dynamic clamp experiments also showed fairly consistent firing responses and clear differences in spike timing reliability under low vs. high frequency inputs. This was not surprising, as the membrane time constant of such developing neurons remains high (cca. 100 ms) while the maximum spiking frequency the neurons reach is well under 20 Hz frequencies. However, the neurons can still participate in network oscillations at lower frequencies (4 Hz or less), and this can contribute to the development of synchronized network activity such as bursting.

According to data published by Odawara et al. (2016), long-term cultivation of h-iPSC-derived cortical neurons cultured on an astrocyte feeder layer resulted in a progressive increase in average firing frequency between week 2 and 34. This indicates a doubling in spike frequency between week 5 and 6 (Odawara *et al*., 2016), with a switch to a more mature phenotype resembling fetal brain tissues after 6 weeks (Sharlow *et al*., 2021). Comparable results were observed regarding maturation timepoints in our h-iPSC-Ns, where a significant increase in neuronal maturation between 4-6 weeks was evident by both the passive and active electrophysiological properties. Our data and previously published studies indicate that a minimum of 4-5 weeks of maturation is needed to attain a fully stabilized electrophysiological phenotype (Odawara *et al*., 2016; Sharlow *et al*., 2021). Comparable outcomes for the active and passive membrane properties of the human neurons were also reported in other studies. For example, (Prè *et al*., 2014; Gunhanlar *et al*., 2018) reported a decline in membrane resistance and membrane time constant, which is consistent with neuronal development over 4-8 weeks Notably, the expression of Na_v_ increased during neuronal maturation, and the membrane became capable of producing action potentials (AP) following depolarization, as observed by Kawaguchi et al. (Kawaguchi *et al*., 2007). The halfwidth of the APs decreased over time, and the amplitude of the AP increased, as also indicated by our data.

It was also interesting that changes in active membrane properties preceded the maturation of cell morphology in time. The initial neurite extension was slowly followed by further branching, so dendritic arborization accelerated only after the 5^th^ week of cultivation, resulting in a more complex cell morphology from the 6^th^ week on. Detailed analyses of morphological maturation of iPSC-Ns also indicated that the neurites of cultured neurons increase in length from DIV 3-9, but both the area of the soma and the length of neurites decreased on subsequent development days (Kang *et al*., 2017). Our morphological analysis revealed a biphasic change in the total dendritic length and process extension between the first 5 and later weeks of maturation, preceding the time-correlated network activity at week 8. The occurrence of the calcium transients highlighted by the synchronicity of the neuronal network additionally increased in a time dependent manner between the 4-8 weeks which is consistent with the development of the electrophysiology parameters (Prè et al., 2014).

Network activity was based on the development and action of abundant fast glutamatergic and GABAergic synaptic connections, both imposing depolarizing action on neurons, similarly to *in vivo* neural development (Dehorter *et al*., 2012). Although a significant change in the expression of GABA_A_ subunits was observed during the 8-week maturation of the cultures, the shift from depolarizing to hyperpolarizing GABA-action did not yet occur. Comparable results of GABA_A_ receptor subunit expression were observed in iCell iPSC-Ns, based on a combination of *alpha* 1/2, *beta* 1/3, and *gamma* 3 subunits (Dage et al., 2014). Yuan et al. (2016) revealed that iCell-Neurons contain a moderate amount of *gamma* 2 mRNA which binds with *alpha* 5 and *beta* 3 subunits to form a unique neuronal subtype distribution when compared to the adult human brain (Yuan *et al*., 2016). This is due to the fact that *alpha* 5 subunits are present in less than 5% of all GABA_A_ receptors in the brain, while *alpha* 1 subunit are present in nearly 60% (Pirker, 2000). In contrast, our developing neurons gradually express the necessary subunits for the GABA_A_ receptors (*alpha* 2, *beta* 3, and *gamma* 2/3 subunits were dominant together with the time-increasing manner of *alpha*1). Interestingly, shifting the expression from early-expressed *NKCC1* to *KCC2* transporters did not occur during the 8-week long cultivation.

Taken together, this study gives us a thematic approach about the neuronal maturation of the human induced pluripotent stem cell derived neurons combining all the prominent key marks of the electrophysiological, morphological, molecular maturation. Our results showed that in order to determine the read-out time of the desired experiments (useful e.g. in testing drug pharmacology), we have to choose the right time of neuronal maturation with a comprehensive approach. Importantly, the presence of electrophysiological responsiveness does not guarantee the appearance of all mature integrative properties. In the same way, morphological phenotypes do not infer the presence of robust network connectivity. Therefore, a given set of these aspects should be used as follow-up standards of the h-iPSC-Ns maturation providing a powerful tool to emphasize the stages of neuronal maturation.

## Acknowledgements

The research is supported by Gedeon-Richter Plc. (Budapest, Hungary) KK/364/2020 grant to K.T. and by ANN_135291 to A.Sz., PD137855 to N.B. and the VEKOP-2.3.3-15-2016-00007 grants from the National Research, Development and Innovation Office, and the ÚNKP-22-3 New National Excellence Program of the Ministry for Culture and Innovation from the source of the National Research, Development and Innovation Fund to BM.M.

## Notes

### Competing Interest Statement

The authors have declared no competing interest.

## References

Ardhanareeswaran, K. et al. (2017) ‘Human induced pluripotent stem cells for modelling neurodevelopmental disorders’, Nature Reviews Neurology, 13(5), pp. 265–278. doi: 10.1038/nrneurol.2017.45.

Arshadi, C. et al. (2021) ‘SNT: a unifying toolbox for quantification of neuronal anatomy’, Nature Methods, 18(4), pp. 374–377. doi: 10.1038/s41592-021-01105-7.

Autar, K. et al. (2022) ‘A functional hiPSC-cortical neuron differentiation and maturation model and its application to neurological disorders’, Stem Cell Reports, 17(1), pp. 96–109. doi: 10.1016/j.stemcr.2021.11.009.

Aversano, S., Caiazza, C. and Caiazzo, M. (2022) ‘Induced pluripotent stem cell-derived and directly reprogrammed neurons to study neurodegenerative diseases: The impact of aging signatures’, Frontiers in Aging Neuroscience, 14(December). doi: 10.3389/fnagi.2022.1069482.

Bean, B. P. (2007) ‘The action potential in mammalian central neurons’, Nature Reviews Neuroscience, 8(6), pp. 451–465. doi: 10.1038/nrn2148.

Berry, B. J. et al. (2019) ‘Advances and current challenges associated with the use of human induced pluripotent stem cells in modeling neurodegenerative disease’, Cells Tissues Organs, 205(5–6), pp. 331–349. doi: 10.1159/000493018.

Dage, J. L. et al. (2014) ‘Pharmacological characterisation of ligand- and voltage-gated ion channels expressed in human iPSC-derived forebrain neurons’, Psychopharmacology, 231(6), pp. 1105–1124. doi: 10.1007/s00213-013-3384-2.

Dehorter, N. et al. (2012) ‘Timing of developmental sequences in different brain structures: Physiological and pathological implications’, European Journal of Neuroscience, 35(12), pp. 1846–1856. doi: 10.1111/j.1460-9568.2012.08152.x.

Deneault, E. et al. (2019) ‘CNTN5 -/+ or EHMT2 -/+ human iPSC-derived neurons from individuals with autism develop hyperactive neuronal networks’, eLife, 8, pp. 1–26. doi: 10.7554/eLife.40092.

Gunhanlar, N. et al. (2018) ‘A simplified protocol for differentiation of electrophysiologically mature neuronal networks from human induced pluripotent stem cells’, Molecular Psychiatry, 23(5), pp. 1336–1344. doi: 10.1038/mp.2017.56.

Hernáth, F., Schlett, K. and Szücs, A. (2019) ‘Alternative classifications of neurons based on physiological properties and synaptic responses, a computational study’, Scientific Reports, 9(1), pp. 1–16. doi: 10.1038/s41598-019-49197-8.

Hutcheon, B., Morley, P. and Poulter, M. O. (2000) ‘2000_Hutcheon&Poulter_Journal-Physiology’, pp. 3–17.

Kang, S. et al. (2017) ‘Characteristic analyses of a neural differentiation model from iPSC-derived neuron according to morphology, physiology, and global gene expression pattern’, Scientific Reports, 7(1), pp. 1–11. doi: 10.1038/s41598-017-12452-x.

Kawaguchi, A. et al. (2007) ‘Enhancement of sodium current in NG108-15 cells during neural differentiation is mainly due to an increase in NaV1.7 expression’, Neurochemical Research, 32(9), pp. 1469–1475. doi: 10.1007/s11064-007-9334-9.

Nagy, J. et al. (2017) ‘Altered neurite morphology and cholinergic function of induced pluripotent stem cell-derived neurons from a patient with Kleefstra syndrome and autism’, Translational Psychiatry, 7(7). doi: 10.1038/TP.2017.144.

Nicholson, M. W. et al. (2022) ‘Utility of iPSC-Derived Cells for Disease Modeling, Drug Development, and Cell Therapy’, Cells, 11(11). doi: 10.3390/cells11111853.

Nowotny, T. et al. (2006) ‘StdpC: A modern dynamic clamp’, Journal of Neuroscience Methods, 158(2), pp. 287–299. doi: 10.1016/j.jneumeth.2006.05.034.

Odawara, A. et al. (2016) ‘Physiological maturation and drug responses of human induced pluripotent stem cell-derived cortical neuronal networks in long-term culture’, Scientific Reports, 6(February), pp. 1–14. doi: 10.1038/srep26181.

Olova, N. et al. (2019) ‘Partial reprogramming induces a steady decline in epigenetic age before loss of somatic identity’, (October 2018). doi: 10.1111/acel.12877.

Pirker, S. (2000) ‘Neuroscience 2000 Pirker’, 101(4), pp. 1–36. Available at: papers2://publication/uuid/8B3931CC-CB6F-4C2F-8CED-BF35D92B9754.

Prè, D. et al. (2014) ‘A time course analysis of the electrophysiological properties of neurons differentiated from human induced Pluripotent Stem Cells (iPSCs)’, PLoS ONE, 9(7). doi: 10.1371/journal.pone.0103418.

Sahlgren Bendtsen, K. M. and Hall, V. J. (2023) ‘The Breakthroughs and Caveats of Using Human Pluripotent Stem Cells in Modeling Alzheimer’s Disease’, Cells, 12(3). doi: 10.3390/cells12030420.

Schindelin, J. et al. (2012) ‘Fiji: An open-source platform for biological-image analysis’, Nature Methods, 9(7), pp. 676–682. doi: 10.1038/nmeth.2019.

Sharlow, E. R. et al. (2021) ‘Extracellular cues accelerate neurogenesis of induced pluripotent stem cell derived neurons’, bioRxiv, p. 2021.08.17.456634. doi: 10.1101/2021.08.17.456634.

Sheng, C. et al. (2018) ‘A stably self-renewing adult blood-derived induced neural stem cell exhibiting patternability and epigenetic rejuvenation’, Nature Communications. doi: 10.1038/s41467-018-06398-5.

Shi, Y. et al. (2017) ‘Induced pluripotent stem cell technology: A decade of progress’, Nature Reviews Drug Discovery, 16(2), pp. 115–130. doi: 10.1038/nrd.2016.245.

Silva, M. C. and Haggarty, S. J. (2020) ‘Human pluripotent stem cell–derived models and drug screening in CNS precision medicine’, Annals of the New York Academy of Sciences, 1471(1), pp. 18–56. doi: 10.1111/nyas.14012.

Szabó, A., Schlett, K. and Szücs, A. (2021) ‘Conventional measures of intrinsic excitability are poor estimators of neuronal activity under realistic synaptic inputs’, PLoS Computational Biology, 17(9), pp. 1–23. doi: 10.1371/journal.pcbi.1009378.

Szücs, A. and Huerta, R. (2015) ‘Differential effects of static and dynamic inputs on neuronal excitability’, Journal of Neurophysiology, 113(1), pp. 232–243. doi: 10.1152/jn.00226.2014.

Takahashi, K. and Yamanaka, S. (2006) ‘Induction of Pluripotent Stem Cells from Mouse Embryonic and Adult Fibroblast Cultures by Defined Factors’, Cell, 126(4), pp. 663–676. doi: 10.1016/J.CELL.2006.07.024.

Wilson, E. S. and Newell-Litwa, K. (2018) ‘Stem cell models of human synapse development and degeneration’, Molecular Biology of the Cell, 29(24), pp. 2913–2921. doi: 10.1091/mbc.E18-04-0222.

Xie, Y. et al. (2018) ‘Reproducible and efficient generation of functionally active neurons from human hiPSCs for preclinical disease modeling’, Stem Cell Research, 26, pp. 84–94. doi: 10.1016/j.scr.2017.12.003.

Yang, G. and Shcheglovitov, A. (2020) ‘Probing disrupted neurodevelopment in autism using human stem cell-derived neurons and organoids: An outlook into future diagnostics and drug development’, Developmental Dynamics, 249(1), pp. 6–33. doi: 10.1002/dvdy.100.

Yuan, N. Y. et al. (2016) ‘Characterization of GABAA receptor ligands with automated patch-clamp using human neurons derived from pluripotent stem cells’, Journal of Pharmacological and Toxicological Methods, 82, pp. 109–114. doi: 10.1016/j.vascn.2016.08.006.

Zhang, K. et al. (2017) ‘High - level extracellular protein production in Bacillus subtilis using an optimized dual - promoter expression system’, Microbial Cell Factories, pp. 1–15. doi: 10.1186/s12934-017-0649-1.

